# Real-Time Three-Dimensional Tracking of Single Vesicles Reveals the Abnormal Motion and Vesicle Pools of Synaptic Vesicles in Neurons of Huntington’s Disease Mice

**DOI:** 10.1101/2021.01.11.426182

**Authors:** Sidong Chen, Hanna Yoo, Chun Hei Li, Chungwon Park, Li Yang Tan, Sangyong Jung, Hyokeun Park

## Abstract

Although defective synaptic transmission was suggested to play a role in neurodegenerative diseases, the dynamics and vesicle pools of synaptic vesicles during neurodegeneration remain elusive. Here, we performed real-time three-dimensional tracking of single synaptic vesicles in cortical neurons from a mouse model of Huntington’s disease (HD). Vesicles in HD neurons had a larger net displacement and radius of gyration compared with wild-type neurons. Vesicles with a high release probability (P_r_) were interspersed with low-P_r_ vesicles in HD neurons, whereas high-P_r_ and low-P_r_ vesicle pools were spatially separated in wild-type neurons. Non-releasing vesicles in HD neurons had an abnormally high prevalence of irregular oscillatory motion. These abnormal dynamics and vesicle pools were rescued by overexpressing Rab11, and the abnormal irregular motion was rescued by jasplakinolide. These results suggest the abnormal dynamics and vesicle pools of synaptic vesicles in the early stages of HD, suggesting a possible pathogenic mechanism of neurodegenerative diseases.

## Introduction

Neuronal communication, an essential process by which neuronal information is encoded in neuronal networks, is mediated by the highly efficient, precise, and tightly regulated release of neurotransmitters from synaptic vesicles in presynaptic terminals (Sudhof and Rizo, 2011); thus, alteration in the properties of synaptic vesicles can have major effects on basal synaptic transmission (Alabi and Tsien, 2012), as well as short-term and long-term synaptic plasticity (Liu, 2003). Although a large body of evidence has accumulated demonstrating the general physiological and morphological properties of synaptic vesicles (Jahn and Fasshauer, 2012; Tao et al., 2018), the study of single synaptic vesicles in real time before, during, and after exocytosis (i.e., fusion) has remained challenging due to both technical and practical limitations. These limitations include the complexity associated with the three-dimensional structure of presynaptic terminals and the minuscule size of synaptic vesicles, which have an average diameter of approximately 40 nm, well below the resolution of conventional light microscopy (Yu et al., 2016). Recently, we developed a method for tracking the three-dimensional position of single synaptic vesicles in real time in cultured hippocampal neurons using quantum dots (QDs) and dual-focus imaging optics, providing localization with an accuracy on the order of tens of nanometers (Park et al., 2012). In addition, we used this approach to examine the relationship between a single synaptic vesicle’s location and its release probability (P_r_) (Park et al., 2012). Although changes in synaptic vesicle dynamics may play a pathogenic role in the early stages of neurodegenerative diseases such as Alzheimer’s (Zhou et al., 2017), Parkinson’s (Kyung et al., 2018), and Huntington’s disease (HD) (Chen et al., 2018), the precise mechanisms that underlie these changes remain largely unknown, due in large part to the relative paucity of studies regarding the motion and release properties of single synaptic vesicles in the context of neurodegeneration.

Huntington’s disease (HD) is a neurodegenerative disorder caused by an increase in CAG repeats in the huntingtin (*HTT*) gene and a corresponding expansion of the polyglutamine (polyQ) tract in the N-terminus of the huntingtin protein (MacDonald et al., 1993). Although the huntingtin protein is expressed essentially throughout the body, medium spiny neurons in the striatum and pyramidal neurons in the cortex are particularly vulnerable to the polyQ expansion in the huntingtin protein (Vonsattel and DiFiglia, 1998). Under physiological conditions, the huntingtin protein is involved in a variety of cellular functions, including the transport of vesicles and organelles (Saudou and Humbert, 2016; Schulte and Littleton, 2011; Yu et al., 2018), transcriptional regulation (Benn et al., 2008; Saudou and Humbert, 2016; Schulte and Littleton, 2011), and cell survival (Ho et al., 2001; Rigamonti et al., 2001). It is therefore reasonable to speculate that the mutant huntingtin protein with an expanded polyglutamine repeat may disrupts these functions, leading to neuronal death in the striatum and cortex of HD patients. Interestingly, the mutant huntingtin protein has been shown to alter the release of neurotransmitters from synaptic vesicles (Cepeda and Levine, 2020; Chen et al., 2018; Joshi et al., 2009; Romero et al., 2008), contributing to the early onset of synaptic dysfunction in the preclinical stages of HD (Milnerwood and Raymond, 2010; Schippling et al., 2009), although the underlying mechanisms are not clearly understood.

Here, we tracked the real-time three-dimensional position of single synaptic vesicles in cortical neurons cultured from an established HD knock-in mouse model and found the abnormal dynamics of single synaptic vesicles in HD neurons. Moreover, synaptic vesicles with a high release probability (P_r_) were interspersed with low-P_r_ vesicles in HD neurons. Besides, non-releasing synaptic vesicles in HD cortical neurons have an abnormally high prevalence of irregular oscillatory motion. The abnormal dynamics and vesicle pools of releasing synaptic vesicles in HD neurons were rescued by overexpressing Rab11 and the abnormal dynamics of non-releasing vesicle were rescued by stabilizing actin filaments with jasplakinolide. Taken together, we have provided the first observation of the abnormal dynamics and vesicle pools of single synaptic vesicles in the early stages of HD.

## Results

### Synaptic Vesicles in HD Neurons Have Abnormal Motion

First, we examined the motion and release of single synaptic vesicles which were loaded with a single streptavidin-coated quantum dot (QD) conjugated to commercially available biotinylated antibodies against the luminal domain of the vesicular protein synaptotagmin-1 (Figure 1A). For these experiments, we used primary cortical neurons cultured from wild-type (WT) mice and heterozygous zQ175 knock-in mice (HD) expressing the human *HTT* exon 1 sequence containing approximately 190 CAG repeats (Menalled et al., 2012); previous studies showed that these mice are a suitable model for studying neurodegeneration in HD patients, particularly with respect to the underlying genetic defect and the disease’s relatively late onset, slow progression, and neuropathology (Chen et al., 2018; Menalled et al., 2012).

**Figure 1.**
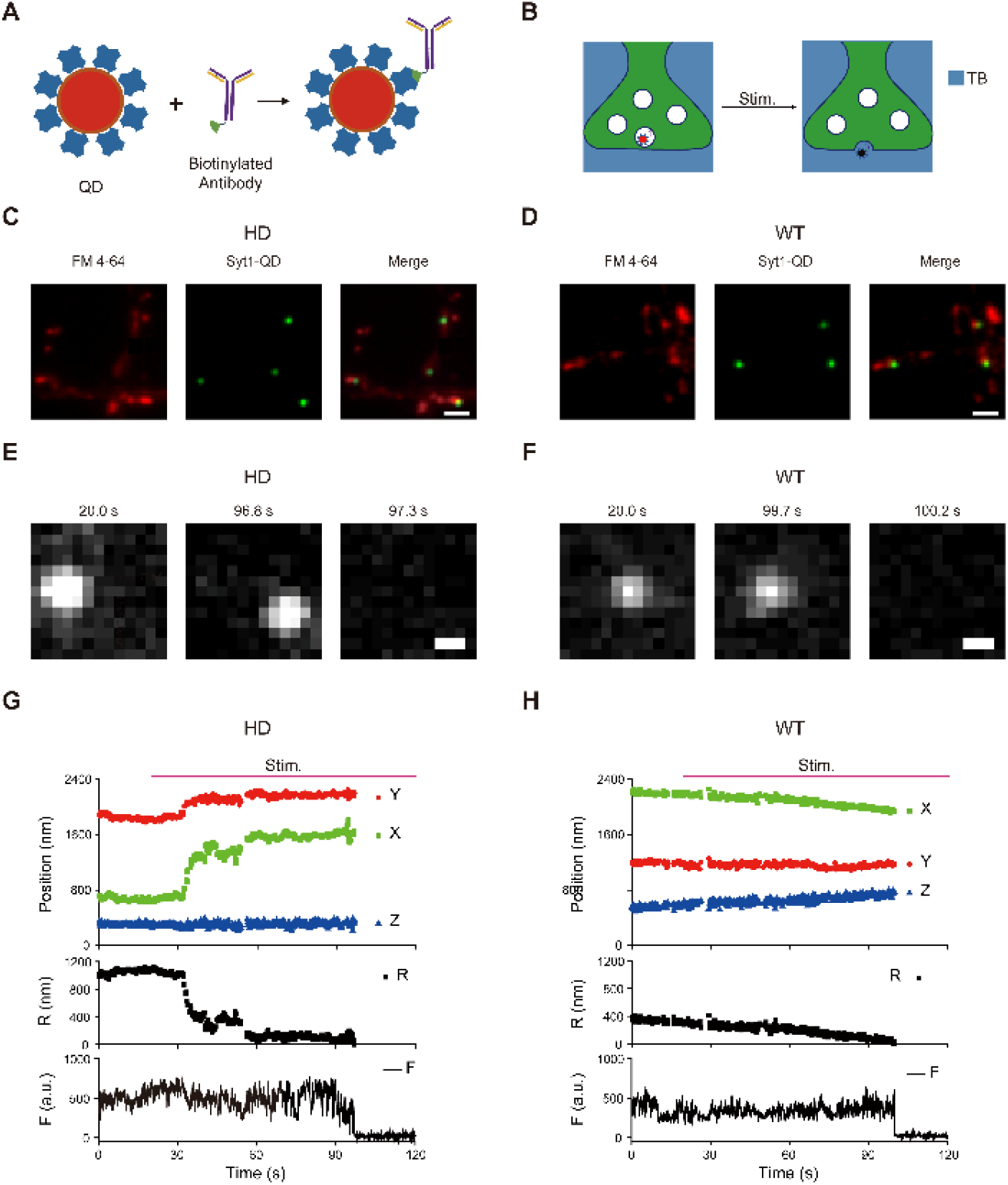
Real-Time Three-Dimensional Tracking of a Single Synaptic Vesicle Loaded with a Single Quantum Dot (QD) in the Presynaptic Terminals of HD and WT Cortical Neurons. (A) Schematic diagram depicting the conjugation strategy. A biotinylated antibody against the luminal domain of synaptotagmin-1 was conjugated to a streptavidin-coated quantum dot (QD) for loading a synaptic vesicle with a single streptavidin-coated QD. (B) Schematic diagram depicting the strategy used to detect exocytosis of QD-loaded synaptic vesicles. After loading, extracellular trypan blue (TB) was used to quench the fluorescence of the QD in the synaptic vesicle during stimulation, thereby indicating the moment of vesicle fusion. (C-D) Colocalization of synaptotagmin-1 (Syt1)-QD-loaded synaptic vesicles (green) and FM 4-64-labeled presynaptic boutons (red) in cultured HD (C) and WT cortical neurons (D). Observed colocalization between the QD and FM 4-64 fluorescence signals indicates that QDs conjugated to antibodies against the luminal domain of Syt1 labeled synaptic vesicles in neurons regardless of genotypes. Scale bar: 2 µm. (E-F) Fluorescence images of the Syt1-QD-loaded vesicle taken at the indicated times in cultured HD (E) and WT cortical neurons (F). Fluorescence images of the QD-loaded synaptic vesicles just before (at 96.8 s in panel E and 99.7 s in panel F) and after exocytosis (at 97.3 s in panel E and 100.2 s in panel F) show near-complete and irreversible quenching of fluorescence regardless of genotypes, indicating exocytosis of QD-loaded synaptic vesicles. Scale bar: 0.5 µm. (G-H) Representative time course of the three-dimensional position, radial distance (R), and QD fluorescence (F) measured in the synaptic vesicle in panel E and F. Where indicated, the neuron was stimulated at 10 Hz stimulation for 120 s. The three-dimensional radial distance was calculated from the momentary position to the fusion site 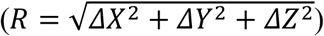.

The exocytosis of single QD-labeled synaptic vesicles during electrical stimulation was reflected by a rapid and irreversible drop in QD fluorescence intensity due to quenching by trypan blue (TB) in the extracellular solution (Figure 1B). Using this loading and quenching protocol, we can track the position of a single synaptic vesicle up until the moment of exocytosis (Park et al., In press; Park et al., 2012). The position of each QD-labeled synaptic vesicle in the *x*-*y* plane was determined to an accuracy on the order of tens of nanometers using FIONA (fluorescence imaging with one-nanometer accuracy) (Park et al., 2007; Yildiz and Selvin, 2005), and the position along the *z*-axis was also determined to an accuracy on the order of tens of nanometers using dual-focusing imaging optics (Park et al., 2012; Watanabe et al., 2007), providing highly accurate three-dimensional trajectories in real time. In order to confirm the labeling of synaptic vesicles with QD-conjugated to biotinylated antibodies against synaptotagmin-1, we used FM 4-64 to label spontaneously released synaptic vesicles and observed colocalization between the QD and FM 4-64 fluorescence signals (Figure 1C - D), which indicates that QDs conjugated to antibodies against the luminal domain of synaptotagmin-1 labeled synaptic vesicles in neurons regardless of genotypes.

Fluorescence images of the QD-loaded synaptic vesicles just before and after exocytosis regardless of genotypes show near-complete and irreversible quenching of QD fluorescence (Figure 1E-F), indicating exocytosis of QD-loaded synaptic vesicles. In addition to the three-dimensional position of QD-loaded synaptic vesicles, we analyzed the radial distance (*R*) from the momentary position to the fusion site (calculated using the equation 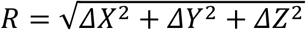, expressed in nm), and relative fluorescence intensity (F) in HD and WT neurons (Figure 1G - H). Fluorescence traces reveal a sharp and irreversible loss of fluorescence at around 97 s (Figure 1G) or 100 s (Figure 1H) caused by exposure of the QDs to trypan blue in the external solution, indicating exocytosis of QD-loaded synaptic vesicles. These traces illustrate that a single synaptic vesicle can be loaded with a single QD by endocytosis and can be successfully tracked in real time until exocytosis in three dimensions during electrical stimulation regardless of genotype. Interestingly, we found that the vesicle in the HD neuron was highly mobile prior to fusion (Figure 1G), whereas the vesicle in the WT neuron was relatively stationary (Figure 1H).

To study the dynamics of releasing synaptic vesicles, we calculated the net displacement between the initial position and the fusion site of individual synaptic vesicles during electrical stimuli at 10 Hz. The three-dimensional net displacement of each vesicle was calculated as the Pythagorean displacement using our real-time three-dimensional traces of single QD-loaded synaptic vesicles. We found that the net displacement of synaptic vesicles in HD neurons was significantly larger compared with synaptic vesicles in WT neurons (242 ± 24.4 nm (average ± standard error of the mean (SEM)) (n = 65 vesicles, N = 14 experiments) vs.170 ± 17.3 nm (n = 80, N = 14 experiments), respectively; *p*=0.0049, Kolmogorov–Smirnov (K-S) test)(Figure 2A). We also estimated the volumetric space in which each synaptic vesicle traveled prior to exocytosis by calculating the three-dimensional radius of gyration (*R*_*g*_) before fusion using the following equation (Qin et al., 2019):

**Figure 2.**
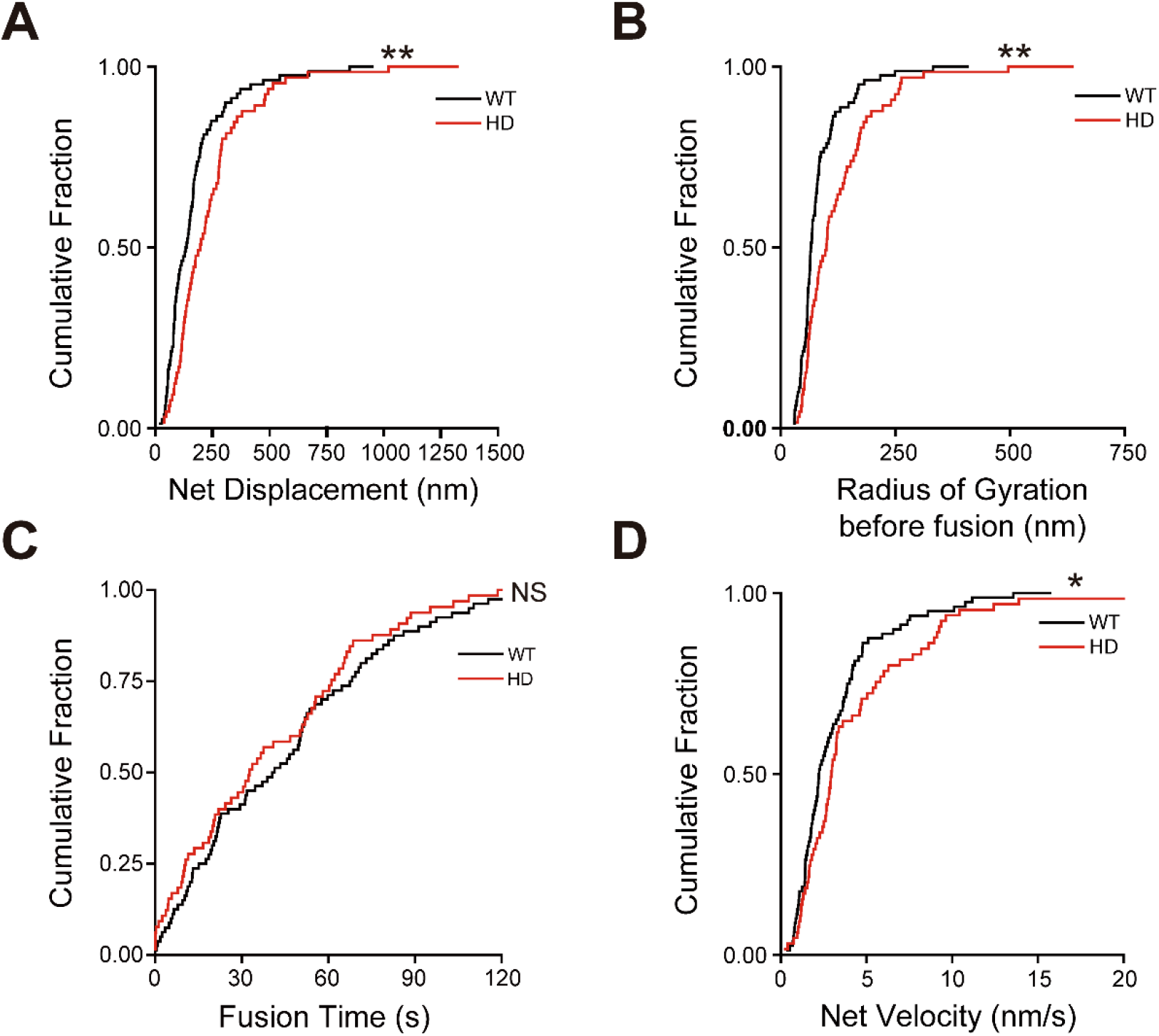
Synaptic Vesicles in HD Cortical Neurons Have Abnormal Motion. (A-D) Cumulative distribution of the net displacement (A), radius of gyration before fusion (B), fusion time (C) and velocity (D) of releasing synaptic vesicles in WT and HD neurons. The net displacement, radius of gyration before fusion and velocity of releasing synaptic vesicles in WT (n = 80 vesicles) were significantly different from those in HD (n = 65 vesicles) neurons. ^*^*p*<0.05, ^**^*p*<0.01 and NS, not significant (Kolmogorov–Smirnov (K-S) test).

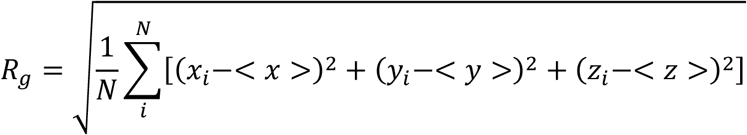

in which *N* is the total number of time steps, *x*_*i*_, *y*_*i*_, and *z*_*i*_ are the projections in the *x-, y-*, and *z-*axes, respectively, at time step *i*, and *<x>, <y>*, and *<z>* are the average positions in each axis. As shown in Figure 2B, the radius of gyration before fusion of synaptic vesicles in HD neurons was significantly larger compared to those in WT neurons (125 ± 11.7 nm vs. 84 ± 6.5 nm, respectively; *p*=0.0023, K-S test)(Figure 2B). Although the fusion time of synaptic vesicles, which is defined as interval between the start of stimulation and fusion, was smaller for HD neuron compared with WT neurons (39.9 ± 3.93 s vs. 45.2 ± 3.83 s, respectively)(Figure 2C), the difference was not statistically significant (*p*=0.87, K-S test). Moreover, synaptic vesicles in HD neurons had higher velocity compared with WT neurons (4.8 ± 0.78 nm/s vs. 2.9 ± 0.25 nm/s, respectively; *p*=0.033, K-S test) (Figure 2D). Taken together, these results indicate that synaptic vesicles in HD neurons are more mobile compared with WT neurons prior to exocytosis, which raise the possibility of an abnormal synaptic vesicle pools in HD neurons.

### Synaptic Vesicles in HD Neurons Have Abnormal Synaptic Vesicle Pools

We previously reported that the vesicle’s release probability (P_r_) is closely related to its synaptic location in rat hippocampal neurons; specifically, we found that synaptic vesicles close to their fusion sites have a higher P_r_ compared with vesicles located relatively far from their fusion sites (Park et al., 2012). Thus, some synaptic vesicles located close to their fusion sites are docked to the presynaptic membrane and primed to rapidly release their contents upon Ca^2+^ influx.

Consistent with these findings, we found that synaptic vesicles with high P_r_ (defined as fusion occurring within 20 s after the onset of stimulation, early SV) are located closer to their fusion sites compared with vesicles with low P_r_ (defined as fusion occurring after 50 s after the onset of stimulation, late SV) in WT neurons (mean net displacement, 111 ± 20.6 (n = 24) nm vs. 192 ± 27.5 (n = 34) nm, respectively; *p*=0.0026, K-S test))(Figure 3A), which indicates that high-P_r_ and low-P_r_ vesicle pools were spatially separated in WT neurons. In contrast, we found no difference in net displacement between high-P_r_ vesicles and low-P_r_ vesicles in HD neurons (222 ± 53.6 nm (n = 22)) vs. 225 ± 29.7 (n = 26) nm respectively; *p*=0.88, K-S test)) (Figure 3B), indicating that high-P_r_ vesicles were interspersed with low-P_r_ vesicles in HD neurons. These results suggest an abnormal relationship between vesicle release probability and location in HD cortical neurons.

**Figure 3.**
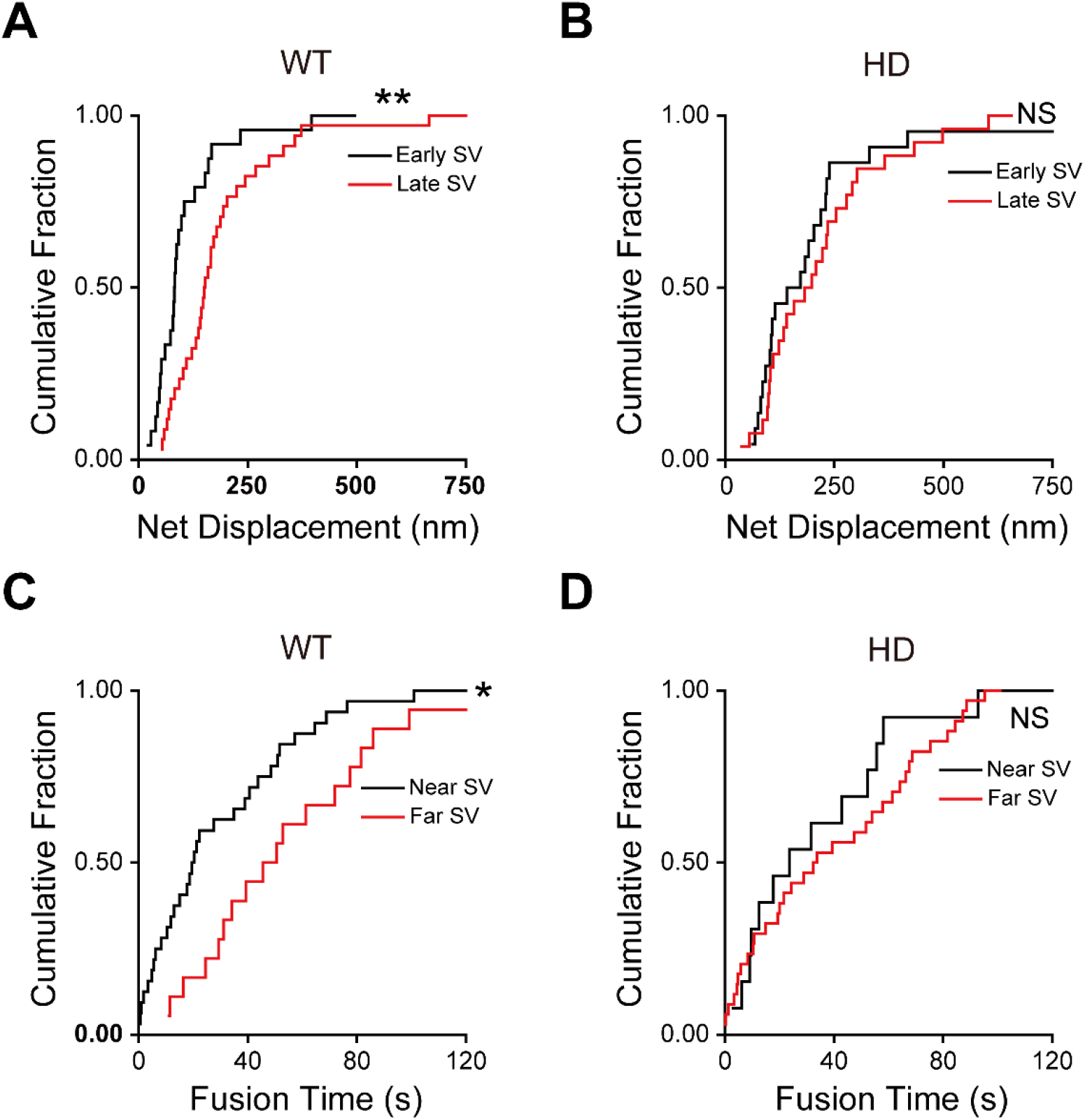
Synaptic Vesicles in HD Neurons Have an Abnormal Relationship between Release Probability (P_r_) and Location. (A-B) Cumulative distribution of the net displacement of early-releasing and late-releasing synaptic vesicles (defined as fusion <20 s or >50 s, respectively, after the start of stimulation) in WT (early SV, n = 24; late SV, n = 34) (A) and HD neurons (early SV, n = 22; late SV, n = 26) (B). (C-D) Cumulative distribution of the fusion time between the onset of stimulation and the fusion of synaptic vesicles initially located near or far from their fusion sites (defined as a net displacement <100 nm or >200 nm, respectively) in WT (near SV, n = 32; far SV, n = 18) (C) and HD neurons (near SV, n = 13; far SV, n = 34) (D). ^*^*p*<0.05,^**^*p*<0.01, and NS, not significant (K-S test).

Next, we examined the relationship between the vesicle’s initial location relative to the fusion site and fusion time. In WT neurons, we found that synaptic vesicles initially located relatively close to their fusion sites (<100 nm (near SV)) had significantly shorter fusion time compared with synaptic vesicles initially located far away from their fusion sites (>200 nm (far SV)) (mean fusion time, 30.2 ± 4.98 s (n = 32) vs. 56.0 ± 8.06 s (n = 18), respectively; *p*=0.030, K-S test) (Figure 3C); this finding is similar to our previous results obtained in rat hippocampal neurons (Park et al., 2012) and supports the close relationship between synaptic vesicle location and release probability. In contrast, we found no such difference in the fusion time for synaptic vesicles close and far from their fusion sites in HD neurons (36.6 ± 9.32 s (n = 13) vs. 40.7 ± 5.38 s (n = 34), respectively; *p*=0.471, K-S test) (Figure 3D). Furthermore, the net displacement of synaptic vesicles was highly correlated with fusion time in WT neurons (Pearson’s r = 0.563, Figure S1A) compared with HD neurons (Pearson’s r = 0.366, Figure S1B). These results support an abnormal relationship between the vesicle’s location and release probability in HD neurons.

Furthermore, we determined whether recycled synaptic vesicles in the readily releasable pool (RRP) relocate back to close to their fusion sites following full-collapse fusion and retrieval from the presynaptic membrane. We tracked the location of the RRP vesicles labeled by applying 10 electrical stimuli at 10 Hz; for comparison, we also tracked the location of vesicles in the total recycling pool (TRP) labeled by applying 1200 electrical stimuli at 10 Hz. We found that in WT neurons, the synaptic vesicles in the RRP were significantly closer to their fusion sites compared to vesicles in the TRP (105 ± 8.0 nm (n = 28) vs. 170 ± 17.3 nm (n = 80) (*p*=0.030, K-S test) (Figure S2A), similar to previous findings in rat hippocampal neurons (Park et al., 2012; Schikorski and Stevens, 2001). In contrast, we found no difference in net displacement between RRP and TRP vesicles in HD neurons (253 ± 40.2 nm (n = 31) vs. 242± 24.4 nm (n = 65), respectively; *p*=0.770, K-S test) (Figure S2B), suggesting that after fusion, RRP vesicles in HD neurons is not relocated close to their fusion site. These results support the notion that synaptic vesicle pools are fundamentally abnormal in HD neurons.

### Non-Releasing Synaptic Vesicles Have Abnormal Oscillatory Motion in the Presynaptic Terminals of HD Neurons

To further examine the abnormal motion of synaptic vesicles in presynaptic terminals of HD neurons, we measured the motion of synaptic vesicles that failed to undergo fusion during the entire electrical stimulation period (i.e., non-releasing vesicles). Interestingly, we found that non-releasing synaptic vesicles in WT neurons generally showed directed (i.e., unidirectional) movement (Figure 4B); in contrast, non-releasing synaptic vesicles in HD neurons often displayed abnormal irregular oscillatory motion, which was characterized by moving back and forth (i.e., bi-directional)(Figure 4A). To quantify this effect, we defined a vesicle with oscillatory motion as having a ratio between the radius of gyration and its net displacement >0.75; considering the size of the average presynaptic terminal, we focused on vesicles with a net displacement <2 μm. We found that HD neurons contained an abnormally large percentage of vesicles with irregular oscillatory motion compared with WT neurons (31 ± 4.0% (N = 14 experiments) vs. 10 ± 2.5% (N = 14), respectively; *p*=0.0001, independent Student’s *t-*test)(Figure 4C). Moreover, this abnormally high incidence of irregular oscillatory motion among non-releasing vesicles in HD neurons compared with non-releasing vesicles in WT neurons translated to a significantly larger travel distance (100 ± 7.0 μm (n = 47 vesicles) vs. 71 ± 4.4 μm (n = 43), respectively; *p*=0.0006, K-S test) (Figure 4D), a significantly larger radius of gyration (216 ± 28.7 nm vs. 119 ± 13.3 nm, respectively; *p*=0.0041, K-S test) (Figure 4E), and a significantly higher instantaneous speed compared with WT vesicles (0.78 ± 0.053 μm/s vs. 0.57 ± 0.028 μm/s, respectively; *p*=0.032, K-S test) (Figure 4F). Taken together, these results support the notion that non-releasing synaptic vesicles in HD neurons have abnormal irregular motion within presynaptic terminals.

**Figure 4.**
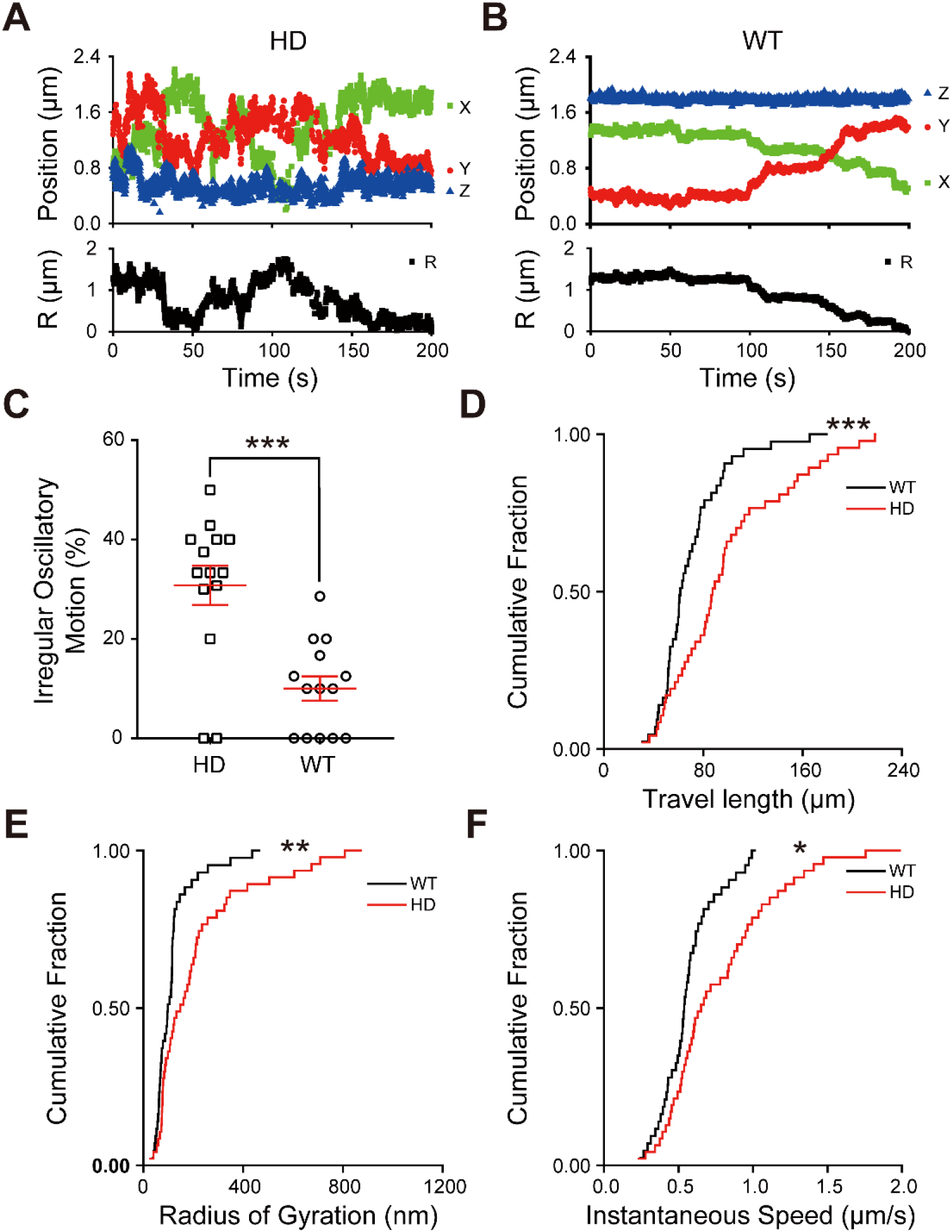
Non-Releasing Synaptic Vesicles in HD Neurons Have Abnormal Irregular Oscillatory Motion. (A-B) Representative time course of the three-dimensional position and radial distance (R) of a single non-releasing synaptic vesicle in a HD (A) and WT neuron (B). (C) Summary of the percentage of synaptic vesicles with irregular oscillatory motion in WT (N = 14 experiments) and HD (N = 14 experiments) neurons. ^***^*p*<0.001 (independent Student’s *t*-test). (D-F) Cumulative distribution of the travel length (D), radius of gyration (E), and instantaneous speed (F) of non-releasing synaptic vesicles in WT (n = 43 vesicles) and HD (n = 47 vesicles) neurons. ^*^*p*<0.05, ^**^*p*<0.01, and ^***^*p*<0.001 (K-S) test).

### Overexpressing Rab11 Rescues the Abnormal Dynamics and Vesicle Pools of Releasing Synaptic Vesicles in HD Neurons

The Ras-related small GTPase Rab11 is present on synaptic vesicles (Sudhof, 2004) and plays a role in vesicle recycling (Kokotos et al., 2018) and endosomal recycling (Ullrich et al., 1996). Moreover, Rab11 has been implicated in several neurodegenerative diseases, including Parkinson’s disease (Breda et al., 2015) and HD (Kiral et al., 2018). For example, reduced Rab11 activity has been reported to impair the formation of synaptic vesicles from recycling endosomes in HD (Li et al., 2009a; Li et al., 2009b). In addition, decreased Rab11 expression was reported in brain lysates obtained from R6/2 mice, a transgenic mouse model with strong phenotypic features associated with HD (Richards et al., 2011). Given these results, we speculated that Rab11 expression may be reduced in cortical neurons in our heterozygotic zQ175 HD mice. Western blot analysis revealed 10% decrease in Rab11 protein levels in the cortical brain lysates of 8-month-old HD mice though the difference was not statistically significant (N = 3 pairs of mice, *p=0*.*10*, independent Student’s *t*-test)(Figure S3). Given the lower expression level of Rab11 and haploinsufficiency of the wild-type huntingtin protein in HD heterozygotic zQ175 cortex, we hypothesized that the abnormal dynamics and vesicle pools of synaptic vesicles in HD neurons may be associated with a functional interaction between the mutant huntingtin protein and Rab11. In support of this hypothesis, we found that Rab11-GFP expressed in HD neurons co-localized with QD-labeled synaptic vesicles (Figure 5A). Moreover, overexpressing Rab11 in HD neurons significantly reduced the net displacement (*p*=0.0016, Figure 5B), radius of gyration before fusion (*p*=0.022, Figure 5C), and velocity (*p*=0.0057, Figure 5D) of releasing synaptic vesicles in HD neurons. However, overexpressing Rab11 did not significantly alter these dynamic properties of releasing synaptic vesicles in WT neurons (Figure 5B-D). Detailed comparison of single synaptic vesicles in WT and HD neurons with expressing Rab11 or an empty vector are summarized in Supplementary Table 1. Two-way ANOVA analyses indicated that the effect of overexpressing Rab11 was significant on the net displacement (*p*=0.011), radius of gyration before fusion (*p<0*.*0001*), and velocity (*p*=0.022) of releasing synaptic vesicles in HD neurons.

**Figure 5.**
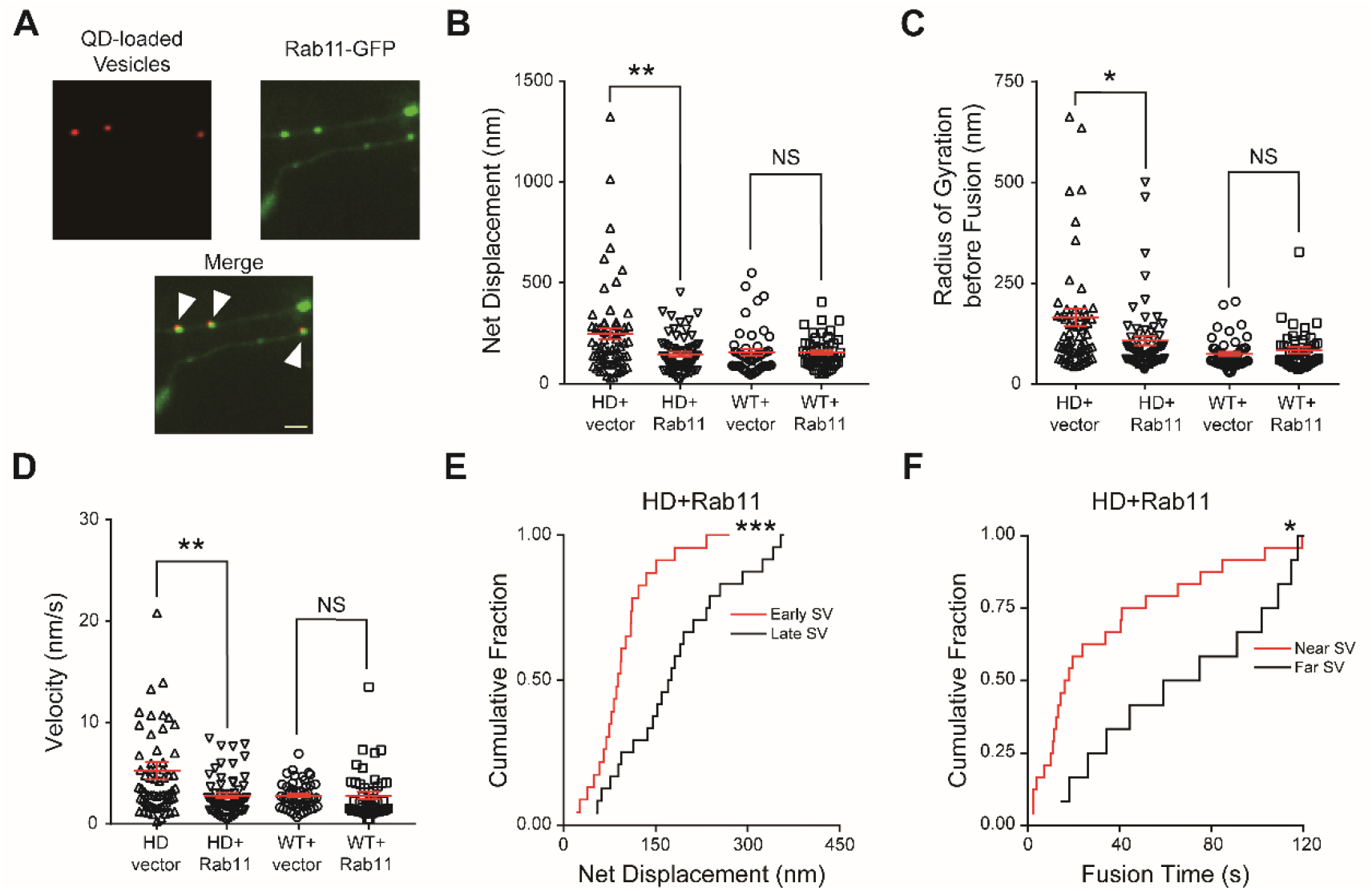
Overexpressing Rab11 Rescues the Abnormal Dynamics and Relationship of Releasing Synaptic Vesicles in HD Neurons. (A) Representative images of QD-loaded vesicles (red) and Rab11-GFP (green) in HD neurons. Arrowheads indicate co-localization between QDs and Rab11-GFP. Scale bar: 2μm. (B-D) Summary of the net displacement (B), radius of gyration before fusion (C), and velocity (D) of releasing vesicles in WT and HD neurons transfected with an empty vector or a vector expressing Rab11-GFP. Overexpressing Rab11 reduced the net displacement (B), radius of gyration before fusion (C), and velocity (D) of releasing synaptic vesicles in HD neurons (empty vector, n = 64 vesicles; Rab11, n = 62 vesicles) to the similar level as WT neurons. In contrast, overexpressing Rab11 did not affect the net displacement (B), radius of gyration (C), and velocity (D) of releasing synaptic vesicles in WT neurons (empty vector, n = 49; Rab11, n = 46). ^*^*p*<0.05, ^**^*p*<0.01 and NS, not significant (independent Student’s *t-*test). (E) Cumulative distribution of the net displacement of early-releasing (n = 23) and late-releasing (n = 24) synaptic vesicles in HD neurons overexpressing Rab11. (F) Cumulative distribution of fusion time of vesicles located near (n = 24) and far (n= 12) from their release sites in HD neurons overexpressing Rab11. ^*^*p*<0.05, ^**^*p*<0.01,^***^*p*<0.001, and NS, not significant (K-S test).

We also tested whether overexpressing Rab11 in HD neurons could rescue the abnormal relationship between the vesicle’s location and its release probability. We found that early-releasing vesicles had significantly smaller net displacement compared with late-releasing vesicles in the overexpression of Rab11 in HD neurons (100 ± 11.4 nm (n = 23, N = 10 experiments) vs. 186 ± 18.7 nm (n = 24, N = 10 experiments), respectively; *p*=0.0008, K-S test)(Figure 4E), similar to our results in rat hippocampal neurons (Park et al., 2012) and our results in WT mouse cortical neurons shown in Figure 3A. In contrast, overexpressing Rab11 had no effect on net displacement in WT neurons (Figure S4A and S4B). We also found that overexpressing Rab11 in HD neurons rescued the abnormal relationship between the vesicle’s initial location and fusion time; specifically, synaptic vesicles located within 100 nm of their fusion sites had significantly shorter fusion time compared with vesicles located >200 nm from their fusion sites in the overexpression of Rab11 in HD neurons (35.1 ± 7.35 s (n = 24) vs. 72.9 ± 11.88 s (n = 12), respectively; *p*=0.037, K-S test)(Figure 5F). In contrast, overexpressing Rab11 had no significant effect in WT neurons (Figure S4C and S4D). Taken together, these results indicate that overexpressing Rab11 can rescue the abnormal relationship between the vesicles’ initial location and their release probability. Interestingly, we found that overexpressing Rab11 did not affect the abnormal oscillatory motion (Figure S5A), net displacement (Figure S5B), and radius of gyration (Figure S5C) of non-releasing vesicles in HD neurons significantly.

### Stabilizing Actin Filaments Rescues the Abnormal Dynamics of Non-Releasing Synaptic Vesicles in the Presynaptic Terminals of HD Neurons

Previous studies showed that the huntingtin protein associates with actin filaments (Angeli et al., 2010; Tousley et al., 2019), possibly mediating the transport of synaptic vesicles to presynaptic terminals (Gramlich and Klyachko, 2017). We therefore hypothesized that stabilizing actin filaments might rescue the abnormal motion of non-releasing vesicles in the presynaptic terminals of HD neurons. Consistent with this hypothesis, we found that treating HD neurons for 10 min with 5 μM jasplakinolide (JKL), which promotes actin polymerization and stabilizes actin filaments (Bae et al., 2012), significantly reduced the irregular oscillatory motion (*p*=0.0003, N = 10 experiments, Figure 6A), travel length (*p*=0.0021, Figure 6B), radius of gyration (*p*=0.0059, Figure 6C), and instantaneous speed (p<0.0001, Figure 6D) of non-releasing synaptic vesicles. In contrast, treating WT neurons with jasplakinolide did not significantly change the irregular oscillatory motion (Figure 6A), travel length (Figure 6B), radius of gyration (Figure 6C), and instantaneous speed (Figure 6D) of non-releasing vesicles. Detailed comparison of non-releasing synaptic vesicle in HD and WT neurons with treating with jasplakinolide or DMSO is summarized in Supplementary Table 2. Two-way ANOVA analyses indicated that treating with jasplakinolide significantly affected the irregular oscillatory motion (*p*=0.0033), travel length (*p*=0.0038), radius of gyration (*p*=0.012), and instantaneous speed (*p*<0.0001) of non-releasing synaptic vesicles in HD neurons. In contrast, jasplakinolide did not have any significant effect on the net displacement of releasing vesicles in HD neurons (Figure S6A) and failed to rescue the abnormal relationship between vesicles’ location and their release probability (Figure S6B and S6C). Thus, jasplakinolide specifically affects the behavior of non-releasing vesicles, but does not affect releasing vesicles in HD neurons.

**Figure 6.**
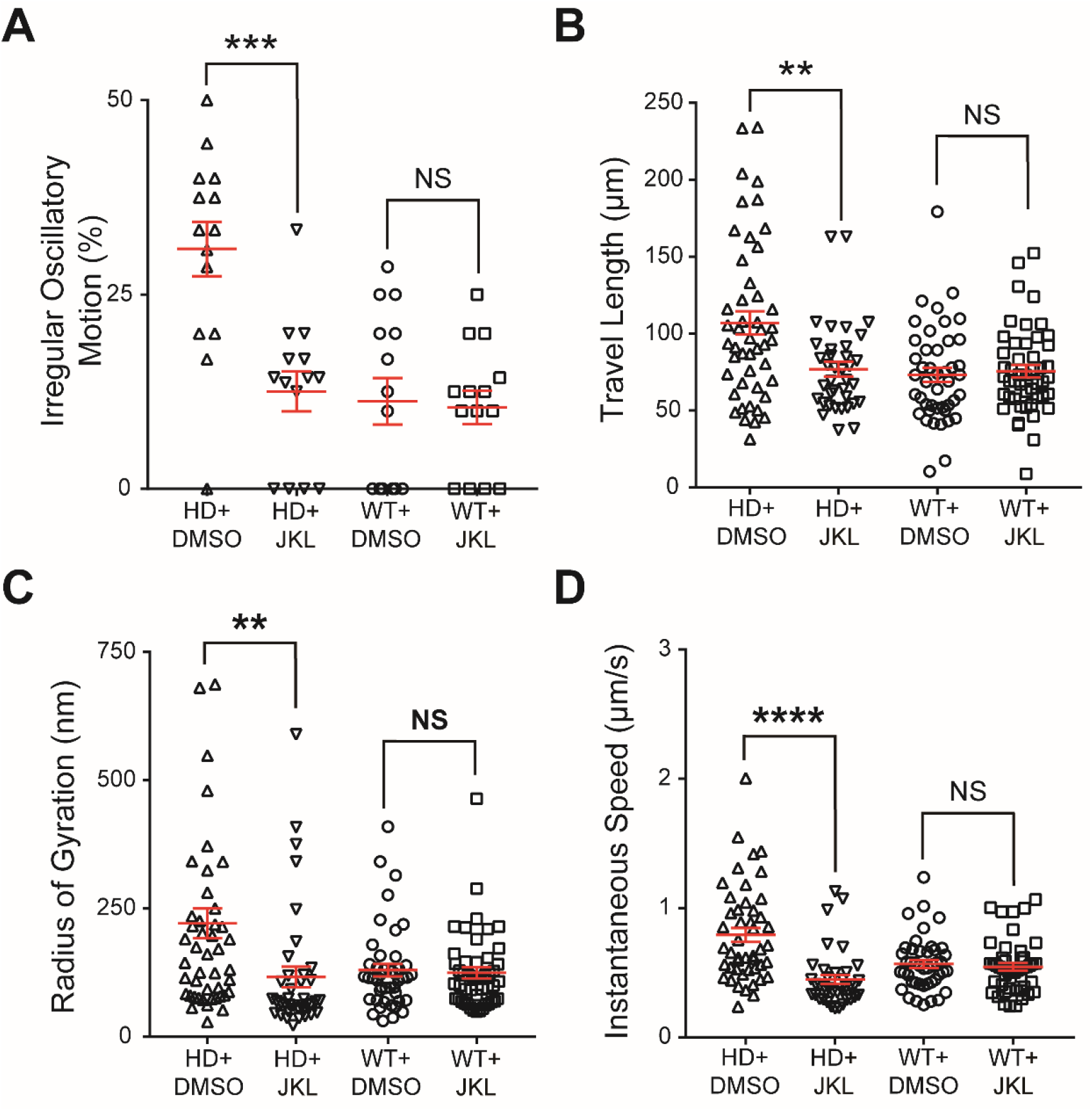
Stabilizing Actin Filaments Rescues the Abnormal Motion of Non-Releasing Synaptic Vesicles in HD Neurons. (A-D) Summary of abnormal irregular oscillatory motion (A), the travel length (B), radius of gyration (C), and instantaneous speed (D) of non-releasing synaptic vesicles in WT and HD neurons treated with DMSO or 5 μM jasplakinolide (JKL). Treating with jasplakinolide reduced the percentage of irregular oscillatory motion (N = 14 experiments for every group) (A), travel length (B), radius of gyration (C), and instantaneous speed (D) of non-releasing synaptic vesicles in HD neurons (DMSO, n = 47; JKL, n = 37) to the similar level as WT neurons. In contrast, jasplakinolide treatment did not affect the percentage of irregular oscillatory motion (A), travel length (B), radius of gyration (C), and instantaneous speed (D) in WT neurons (DMSO, n = 44; JKL, n = 48). ^**^*p*<0.01, ^***^*p*<0.001,^****>*^*p*<0.0001 and NS, not significant (independent Student’s *t-*test).

## Discussion

Although synaptic dysfunction has been suggested to play an important pathogenic role in HD (Chen et al., 2018; Li et al., 2003b; Sepers and Raymond, 2014; Yu et al., 2018), whether the motion and/or vesicle pools of synaptic vesicles is disrupted in HD— particularly with respect to the dynamics of single synaptic vesicles—has not been investigated. To address these questions, we tracked the real-time three-dimensional position of single synaptic vesicles in HD cortical neurons and found the abnormal dynamics and vesicle pools of releasing synaptic vesicles in HD neurons. Moreover, we found that non-releasing synaptic vesicles in the presynaptic terminals of HD neurons have an abnormally high prevalence of irregular oscillatory motion, increasing their travel length and radius of gyration. Importantly, the abnormal dynamics and vesicle pools of releasing synaptic vesicles and the abnormal motion of non-releasing vesicles in HD neurons were rescued by overexpressing Rab11 and stabilizing actin filaments with jasplakinolide, respectively.

Our results suggest a change in neurotransmitter release in HD neurons, which is consistent with previous reports of altered neurotransmitter release in the presynaptic terminals in several HD models (Chen et al., 2018; Joshi et al., 2009; Romero et al., 2008). Any change in synaptic vesicle release will contribute to impaired synaptic transmission in HD, as efficient synaptic transmission requires the accurate, reliable, and precisely timed release of neurotransmitters from synaptic vesicles. In this respect, impaired synaptic transmission may serve as an early pathogenic driver in HD (Milnerwood and Raymond, 2010). Interestingly, we found that treating HD neurons with jasplakinolide (an actin filament stabilizer) rescued the abnormal motion of non-releasing vesicles, suggesting that the mutant huntingtin protein may affect filamentous actin (F-actin) in the presynaptic terminals. Our finding that jasplakinolide does not affect the vesicle pools of releasing vesicles suggests that F-actin alone does not play a role in separating early-releasing vesicles from late-releasing vesicles. On the other hand, we cannot rule out F-actin’s other roles in the recycling of synaptic vesicles, particularly when a synaptic vesicle is internalized by “pinching off” from the presynaptic membrane via an endocytic pit (Wu et al., 2016). However, the irregular oscillatory motion in the presynaptic terminals likely reflects a possible defect in actin-based motility due to actin depolymerization in HD neurons because myosin motor proteins such as myosin II and myosin V have been shown to drive the movement of synaptic vesicles along actin filaments over relatively short distances in the presynaptic terminals (Peng et al., 2012).

Despite our finding that 8-month-old heterozygous zQ175 mice displayed 10% decrease in expression level of Rab11, we found that overexpressing Rab11 rescued the abnormal motion of releasing synaptic vesicles. Interestingly, reduced Rab11 expression was previously reported in HD neurons (Richards et al., 2011), and overexpressing Rab11 in a *Drosophila* HD model rescued the neuronal phenotype (Steinert et al., 2012). Studies have also found that the huntingtin protein facilitates the nucleotide exchange activity of Rab11, leading to Rab11 activation in cortical neurons (Li et al., 2008). In contrast, the mutant huntingtin protein has been shown to interfere with Rab11 activation, reducing endocytic vesicle formation in HD fibroblasts (Li et al., 2009b). Together, these findings suggest that the mutant huntingtin protein may downregulate or activate Rab11 to a lesser extent compared with the wild-type protein, leading to impaired vesicle dynamics in the presynaptic terminals of HD neurons. In this respect, it is interesting to note that overexpressing Rab11 rescue abnormal intermingling between early-releasing vesicles and late-releasing vesicles in HD neurons, suggesting that Rab11 may play an essential role in facilitating the localization of both early-releasing and late-releasing vesicles, and possibly non-releasing vesicles in the reserve pool. Nevertheless, further research is warranted to investigate the detailed mechanism by which Rab11 mediates the localization of synaptic vesicles based on release probability.

Although Rab11 overexpression and jasplakinolide treatment rescued the abnormal dynamics of single synaptic vesicles in our HD mouse model, we cannot rule out the possibility that other proteins and/or processes also contribute to abnormal vesicle dynamics in HD neurons. The huntingtin protein is a large scaffolding protein that interacts with several binding partners (Shirasaki et al., 2012), and disrupting these interactions can result in impaired glutamate release (Li et al., 2003a). Huntingtin-associated protein 1 (HAP1) is a major binding partner and has been shown to regulate the exocytosis of synaptic vesicles and play a role in the actin-based transport of insulin-containing granules in pancreatic beta cells (Mackenzie et al., 2016; Wang et al., 2015). Recently, Rab4, which coordinates vesicle trafficking, was reported to be affected in HD (White et al., 2020). Given that the huntingtin protein can interact with many proteins, some of which may have additional binding partners (Shirasaki et al., 2012), indirect interactions with the mutant huntingtin protein may contribute—at least in part—to the observed abnormal synaptic vesicle dynamics in HD neurons. Future research may therefore provide insight into the role that these associated proteins play with respect to the abnormal dynamics of synaptic vesicles in HD neurons.

Our findings in the early stage of HD are consistent with the disruption in the dynamics and recycling of synaptic vesicles reported in other neurodegenerative diseases, including Alzheimer’s disease (Marsh and Alifragis, 2018) and Parkinson’s disease (Hunn et al., 2015). Pathogenic tau binds to synaptic vesicles and disrupts synaptic vesicles mobility and release (Zhou et al., 2017). Similarly, the aggregation of alpha-synuclein reduces the size of the recycling vesicle pool and impairs synaptic transmission (Nemani et al., 2010; Scott and Roy, 2012). These impairments in synaptic vesicles dynamics and localization are likely mediated by cytoskeletal defects and Ras-associated small GTPases such as Rab11 (Breda et al., 2015; Udayar et al., 2013). Thus, overlap in the mechanisms that underlie different forms of neurodegeneration may be exploited when developing general therapies for neurodegenerative diseases.

In this study, we performed real-time three-dimensional tracking of single synaptic vesicles in cultured cortical neurons and found that synaptic vesicles in presymptomatic HD mice have the abnormal dynamics and vesicle pools of releasing synaptic vesicle, and abnormal irregular oscillatory motion of non-releasing synaptic vesicles. We also found that stabilizing actin filaments rescued the abnormal dynamics of non-releasing synaptic vesicles in HD neurons, whereas overexpressing Rab11 in HD neurons rescued both the dynamics and vesicle pools of releasing vesicles. Together, these results suggest that the abnormal synaptic vesicles dynamics in the presynaptic terminals of HD neurons arise from the disrupted functions of actin filaments and Rab11, leading to impaired synaptic transmission in the early stage of HD. Thus, our results provide new insights into the role that synaptic vesicle dynamics play in the pathogenesis of HD and other neurodegenerative diseases.

## STAR★Methods

### Mice

Mice of zQ175 (a Huntington’s disease (HD) knock-in mouse model) were obtained from Jackson Laboratories and housed in the Animal and Plant Care Facility at the Hong Kong University of Science and Technology. Only heterozygous zQ175 mice were used for breeding. All procedures for mice handling were approved by Department of Health, Government of Hong Kong. All procedures were performed in accordance with approved protocols.

### Primary cortical neuron cultures

Primary cortical neurons were prepared and cultured as described previously (Chen et al., 2018). Heterozygous zQ175 (HD) and WT neurons were collected from the cerebral cortex of postnatal day 0 (P0) heterozygous zQ175 pups and WT littermates. The dissected cortical neurons were digested briefly with papain (LS003127, Worthington Biochemical Corp., Lakewood, NJ, USA) and DNAse (D5025, Sigma-Aldrich). After gentle trituration, the density of neurons was determined, and approximately 10^5^ neurons were plated on 12-mm glass coverslips precoated with poly-D-lysine (P7405, Sigma-Aldrich) in a 24-well plate as described previously (Chen et al., 2018). After 3 days in culture (DIV3), 20 μM 5-fluoro-2’-deoxyuridine (F0503, Sigma-Aldrich) was added to the culture medium to inhibit the proliferation of glial cells (Chen et al., 2018). The neurons were incubated at 37°C in humidified air containing 5% CO_2_, and real-time imaging was performed at DIV14-17.

### Three-dimensional tracking of single QD-loaded synaptic vesicles

Real-time imaging was performed similarly as described previously (Alsina et al., 2017; Chen et al., 2018) using an IX73 inverted microscope (Olympus) equipped with a 100x oil-immersion UPlanSAPO objective (Olympus) and a dual-focus imaging optics system (Park et al., 2012). Real-time three-dimensional nanometer-accuracy tracking of QD-loaded vesicles was performed as previously described (Park et al., 2012). The real-time fluorescence images at two focus planes were captured side-by-side using an iXon Ultra EMCCD camera (Andor Technology Ltd., Belfast, UK) and a custom-made dual-focus imaging optics system. Custom programs written in IDL (Harris Geospatial Solutions, Inc.) were used to calculate the peak intensities and two-dimensional centroids of the QD at two different focal planes (*I*1 and *I*2) by fitting a two-dimensional Gaussian function to the fluorescence images. A PIFOC piezo stage (Physik Instrumente GmbH, Karlsruhe, Germany) was used to generate a calibration curve representing the relationship between the *z*-position and the relative difference in peak intensity (*I*_1_ - *I*_2_)/(*I*_1_ + *I*_2_). The *z*-position was then calculated from the calibration curve by applying the corresponding value of (*I*_1_ - *I*_2_)/(*I*_1_ + *I*_2_).

### Real-time imaging of single QD-labeled synaptic vesicles in cultured neurons

Streptavidin-coated QDs (A10196, Thermo Fisher Scientific) were conjugated to biotinylated Syt1 antibodies (105 103BT, Synaptic Systems) by incubating at room temperature for one hour. The QD-conjugated antibodies were then added to a sample chamber containing a coverslip with attached neurons. Electrical field stimuli (1200 stimuli applied at 10 Hz for 120 s or 10 stimuli applied at 10 Hz for 1 s) were then applied in order to trigger the exocytosis and endocytosis of synaptic vesicles, causing the vesicles to take up the antibodies-conjugated QDs; the stimuli were applied using a platinum parallel electrode connected to an SD9 Grass Stimulator (Grass Technologies). The Grass stimulator, beam shutter, and EMCCD camera were synchronized with a trigger from the camera using a Digidata 1550 interface (Molecular Devices), and Clampex (Molecular Devices) was used to generate the stimulation protocols. After stimulation and an additional 3-min incubation period, the chamber was rinsed extensively for 10 min with artificial cerebrospinal fluid (ACSF) containing (in mM): 120 NaCl, 4 KCl, 2 CaCl_2_, 2 MgCl_2_, 10 D-glucose, and 10 HEPES (300-310 mOsm, pH 7.2-7.4 with NaOH). Prior to the experiment, trypan blue was added to the ACSF at a final concentration of 2 μM. A 405 nm laser (Coherent Inc., USA) was used to excite QDs, and a dichroic mirror (ZT405rdc, Chroma) and an emission filter (ET605/70m, Chroma) were used to capture the fluorescence signals. Real-time imaging experiments were performed at 10 Hz with an exposure time of 0.1 s for 200 s using the frame transfer mode of the EMCCD camera. After collecting a baseline of 20 s, 1200 electrical field stimuli (10 Hz for 120 s) were applied to the neurons, followed by an addition 60 s without stimulation.

### Western blot analysis

Cortical brain tissues were dissected from 4-month-old WT and heterozygous zQ175 mice and lysed using N-PER Neuronal Protein Extraction Reagent (87792, Thermo Fisher Scientific). The lysates were centrifuged for 21 min at 4°C, and the supernatant was boiled in sample buffer containing 120 mM Tris-HCl, 4% SDS, 20% glycerol, 5% μ-mercaptoethanol, and 0.1 mg bromophenol blue. The samples were then resolved using SDS-PAGE and transferred to a PVDF membrane. The membranes were blocked in 5% (w/v) dry milk in TBS-T solution (0.1% Tween-20 in Tris-buffered saline) and then incubated overnight at 4°C in the following primary antibodies diluted in TBS-T containing 3% (w/v) BSA: rabbit anti-Rab11A (715300, Invitrogen) and rabbit anti-β-actin (4967, Cell Signaling Technology). After washing with TBS-T, the membranes were incubated with HRP-conjugated secondary antibodies at room temperature for 1 h, washed in TBS-T, and visualized using Clarity Western ECL Substrate (Bio-Rad Laboratories) with the ChemiDoc imaging system (Bio-Rad Laboratories). ImageJ was used to quantify the protein levels of Rab11 in both WT and HD tissues.

### Overexpression of Rab11-GFP

Lipofectamine 2000 (11668019, Thermo Fisher Scientific) was used to transfect cultured cortical neurons at DIV7 with a construct expressing Rab11-GFP (plasmid #12674, Addgene) or an empty vector. Transfected neurons were used for experiments starting at DIV14, and only synaptic vesicles in positively transfected neurons were analyzed.

### Jasplakinolide treatment

After the addition of trypan blue to the extracellular solution, jasplakinolide (5 μM; J7473, Thermo Fisher Scientific) or vehicle (DMSO) was added to the chamber, and imaging experiments were performed 10 min later.

### Quantification and Statistical Analysis

The fluorescence intensity within a given region of interest (ROI) was analyzed using MetaMorph software (Molecular Devices). Custom programs written in IDL (Harris Geospatial Solutions, Inc.) were used to calculate the peak intensities and two-dimensional centroids. The differences in net displacement, travel length, fusion time, velocity, and radius of gyration between HD and WT vesicles were analyzed using the Kolmogorov–Smirnov test (GraphPad Prism 7). The differences in the fraction of synaptic vesicles with oscillatory motion and synaptic vesicle dynamics, including rescue experiments, were analyzed using an independent Student’s *t-*test. The effects of Rab11 overexpression and jasplakinolide treatment were analyzed using a two-way ANOVA. Differences were considered significant at *p*<0.05. The Pearson’s r and linear regression line of fusion time and net displacement relationship in HD and WT were calculated by OriginPro 9.

## Acknowledgments

We thank Dr. Richard W. Tsien for generous support and helpful discussions, Dr. Sukho Lee for helpful discussions, and Dr. Curtis F. Barrett for critically reading the manuscript. This work was supported by grants from the Research Grants Council of Hong Kong (26101117, 16101518, N_HKUST613/17, and A-HKUST603/17 to H.P.), the Innovation and Technology Commission (ITCPD/17-9 to H.P.) and Joint Council Office (Grant No. BMSI/15-800003-SBIC-OOE to SJ).

## Author contributions

S.C. and H.P. designed the experiments. S.C., H.Y., C.H.L., and C.P. performed the experiments. S.C. and C.P. analyzed the data. S.C., L.Y.T., S.J., and H.P. wrote the manuscript. All authors read and approved the final manuscript.

## Declaration of Interests

The authors have no competing interests to declare.

## Notes

### Competing Interest Statement

The authors have declared no competing interest.

